# pCMLM: Genome Wide Association Study of Body Size Traits in Multiple Regions of Yak Based on the Provided Compressed Mixed Linear Model

**DOI:** 10.1101/2022.09.26.509454

**Authors:** Xinrui Liu, Zhixin Chai, Wei Peng, Yixi Kangzhu, Jincheng Zhong, Jiabo Wang

## Abstract

**Objective:** Yak is a unique large animal species living in the Qinghai-Tibet Plateau and the surrounding Hengduan Mountains, and has evolved several regional variety resources due to the special geographical and ecological environment in which it lives. Therefore, it is of great importance to investigate the genetic composition of body size traits among breeds in multiple regions for yak breeding and production.

**Method:** A genome-wide association analysis was performed on 94 yak individuals (a total of 31 variety resources) for five body size traits (body height, body weight, body length, chest circumference, and circumference of cannon bone). The individuals were clustered following known population habitat. The kinship of grouping individuals was used in the CMLM. This provided compressed mixed linear model was named pCMLM method.

**Result:** Total of 3,584,464 high-quality SNP markers were obtained on 30 chromosomes. Principal component analysis using the whole SNPs do not accurately classify all populations into multiple subpopulations, a result that is not the same as the population habitat. Six SNP loci were identified in the pCMLM-based GWAS with statistically significant correlation with body height, and four candidate genes (*FXYD6, SOHLH2, ADGRB2*, and *OSBPL6*), which in the vicinity of the variant loci, were screened and annotated. Two of these genes, *ADGRB2* and *OSBPL6*, are involved in biological regulatory processes such as body height regulation, adipocyte proliferation and differentiation.

**Conclusion:** Based on the previous population information, the pCMLM can provide more sufficient associated results when the conventional CMLM can not catch optimum clustering groups. The fundamental information for quantitative trait gene localization or candidate gene cloning in the mechanism of yak body size trait formation.

## 1 Introduction

Yak (*Bos grunniens*) is a large livestock species unique to the Qinghai-Tibet Tibetan Plateau and the surrounding Hengduan Mountains, providing a basic resource for the livelihood of plateau farmers and herders **[Ran H, et al. 2022].** There are 12 breeds of yak in China due to the different geographical and climatic environments, ecological conditions, grassland types, feeding levels, breeding levels, and social and economic structures in the main producing areas. It mainly includes Qinghai Plateau yaks, Gannan yaks, Tianzhu White yaks, Bazhou yaks, Zhongdian yaks, Jiulong yaks, Muli yaks, Maiwa yaks, Niangya yaks, Xizang Alpine yaks and Sibu yaks **[Wu J. 2016]**. Among them, Tibet is one of the main yak-producing areas in China, and yaks are distributed in 71 counties in Tibet, accounting for 30% of the total number of yaks in China, and gradually formed many local yak breeds and groups, such as Niangya yak, Pari yak, Sibu yak, Sangsang yak, Sangri yak, Baqing yak, Dingqing yak, Kangbu yak, Jiangda yak, Leiwuqi yak, Gongbujiangda yak have been formed **[Zhang G, et al. 2008]**.

The rich genetic diversity is the material basis for yak to adapt to changes in the external environment and a vital genetic resource for breeding new breeds or strains. Still, the excellent traits of each breed resource are not actively exchanged during the evolutionary process due to the natural and social-ecological environment, natural selection, and artificial selection. Those were making the current situation that the unique excellent traits of regional yak cannot be outflowed. The body size trait of yak is the most critical genetic index and an essential reference index to determine meat production performance and is also one of the most direct breeding selection parameters **[Junior G A O, et al. 2021]**. Some yaks are more aggressive due to mixing with wild blood, and extracting relevant biological traits is more complicated. And nowadays, the technology of extracting animal body sizes traits by image recognition are getting mature **[Zhang A, et al. 2018; Khojastehkey M, et al. 2016]**. And Gomes R A et al. **[Gomes R A, et al. 2016]** used digital images of beef cattle acquired by Microsoft Kinect device to establish model equations for predicting body, carcass weight, and body fat content to estimate relevant traits such as body weight of beef cattle has become a reality, which makes body sizes trait determination easier and faster day by day.

Developing high-throughput genotyping technologies has provided opportunities to identify novel genetic variants associated with economic traits in cattle, where Single Nucleotide Polymorphisms (SNP) widely distributed throughout the genome have become the genetic markers of choice. Genome-wide association study (GWAS) is a common experimental approach to study SNP markers associated with various economic traits in animal production by linking phenotypic and genotypic data and using statistical models to investigate genetic variant loci causally associated with the target trait.

The GWAS approach has successfully revealed genetic determinants associated with disease susceptibility and resistance in humans **[Sherva R, et al. 2011]**, animals **[Psifidi A, et al. 2016]**, and plants **[Patron A, et al. 2016]**. The compress Mixed Linear Model (CMLM) was reported to improve the statistics power of GWAS by using clustering individuals into groups based on kinship among individuals **[Zhang, et al. 2010]**. When the likelihood values of testing model in the CMLM can not catch the optimum clustering group, the individual will be grouped each group with only one individual. This result has been proved by simulation results. However, the previous population clustering was widespread in the animals, especially in the yaks. The application of previous population structure is the key of animal GWAS.

To address this need, we performed GWAS in the yaks population, targeting the incorporation of two important features:1) to detect associated SNPs with yak body size, and 2) to develop an approach for animal GWAS using previous population habitat.

## 2 Material and method

### 2.1 Individual samples and sequencing(Population and Phenotypic Data)

Total 94 yaks were collected from different regions of the Qinghai-Tibet Tibetan Plateau in China, including 17 Tibetan breeds, 4 Qinghai breeds, 4 Sichuan breeds, 2 Gansu breeds, 2 Xinjiang breeds, 1 Yunnan breed, and 4 wild yak data, for a total of 31 yak breed resources (Table 1 and Figure 2A). Detailed information on the resources of each species and their distribution areas are shown in Table S1. The population samples were obtained from the Key Laboratory of Qinghai-Tibetan Plateau Animal Genetic Resource Reservation and Utilization, Sichuan Province, and Ministry of Education, Southwest Minzu University **[Chai Z X, et al. 2020]**. And their sequence files were all obtained from the sequencing results of DNA extracted from blood samples by the Illumina Hiseq 2000 sequencer (Illumina, San Diego, CA, USA). Phenotypic data were measured after adult (more than 2.5 years old) including records of body height (BH), body length (BL), body weight (BW), chest circumference (CC), circumference of cannon bone (CCB).

**Table.1.**
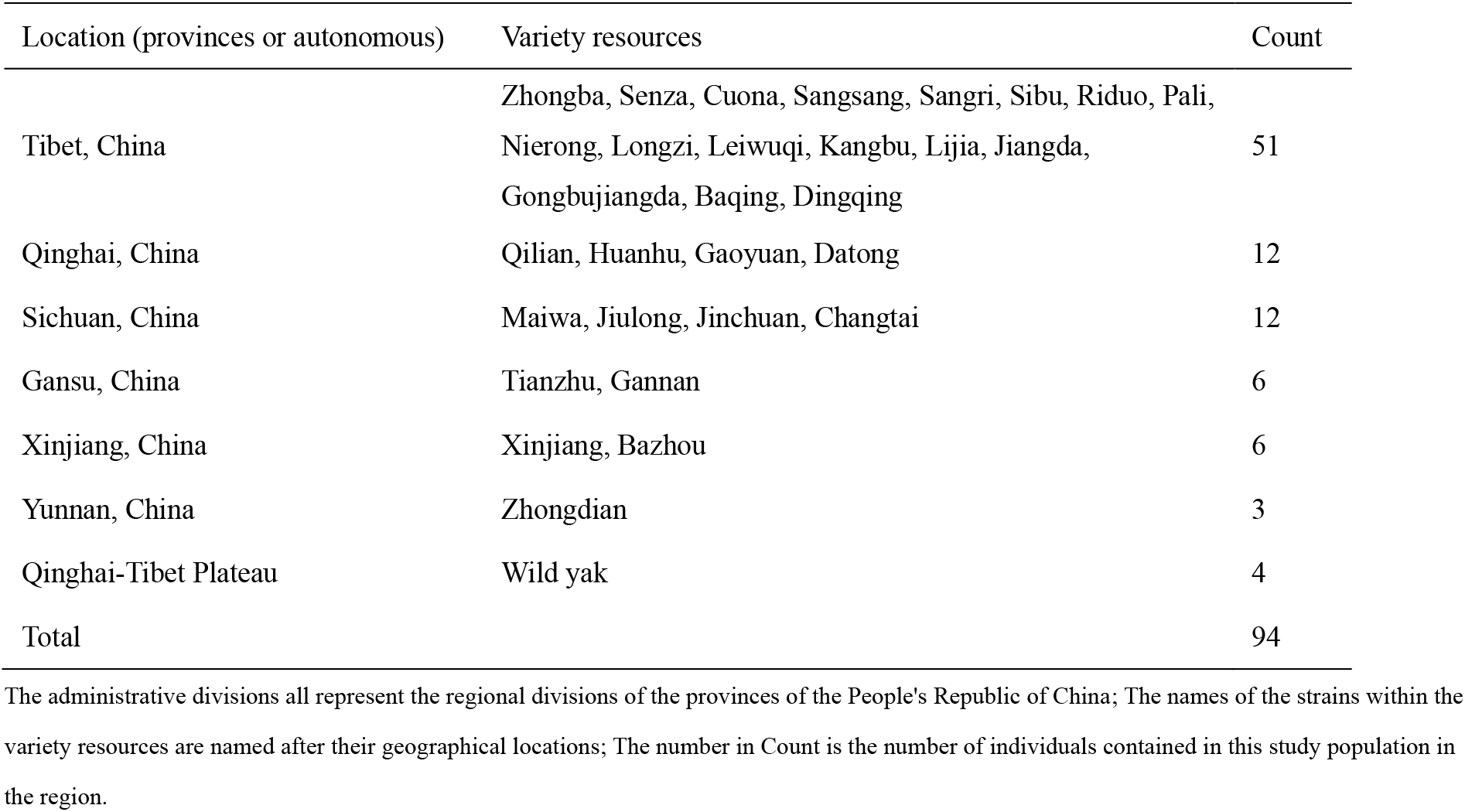
Data distribution of yak samples

| Location (provinces or autonomous) | Variety resources | Count |
| --- | --- | --- |
| Tibet, China | Zhongba, Senza, Cuona, Sangsang, Sangri, Sibu, Riduo, Pali, Nierong, Longzi, Leiwuqi, Kangbu, Lijia, Jiangda, Gongbujiangda, Baqing, Dingqing | 51 |
| Qinghai, China | Qilian, Huanhu, Gaoyuan, Datong | 12 |
| Sichuan, China | Maiwa, Jiulong, Jinchuan, Changtai | 12 |
| Gansu, China | Tianzhu, Gannan | 6 |
| Xinjiang, China | Xinjiang, Bazhou | 6 |
| Yunnan, China | Zhongdian | 3 |
| Qinghai-Tibet Plateau | Wild yak | 4 |
| Total |  | 94 |

The administrative divisions all represent the regional divisions of the provinces of the People's Republic of China; The names of the strains within the variety resources are named after their geographical locations; The number in Count is the number of individuals contained in this study population in the region.

**Table.2.** Descriptive statistics of body measurements traits in yak

| Taxa | Max | Min | Mean | SD | SE | CV |
| --- | --- | --- | --- | --- | --- | --- |
| BH | 205 | 100.7 | 118.2 | 16.7559 | 1.72823 | 14.1805 |
| BL | 240 | 105.4 | 138.6 | 27.3521 | 2.82116 | 19.7283 |
| BW | 821 | 156.1 | 284.9 | 103.982 | 10.7249 | 36.4992 |
| CC | 270 | 140.1 | 168.8 | 23.2648 | 2.39958 | 13.7847 |
| CCB | 22.9 | 10.05 | 17.24 | 2.9005 | 2.39958 | 16.8203 |

Body height (BH, cm), Body length (BL, cm), Body weight (BW, kg), Chest circumference (CC, cm), Circumference of cannon bone (CCB, cm); Maximum (Max), Minimum (Min)Standard deviation (SD), Standard error (SE), Coefficient of variation (CV); The description statistics of all phenotypes are generated by the summary function of R.

### 2.2 Genotyping quality control and filtering

Data filtering was performed using FASTP **[Chen S, et al. 2018]** software(version 0.20.1). Double-end sequencing was spliced using Burrows-Wheeler-Alignment Tool (BWA) **[Li H, et al. 2009]** software (version 0.7.15), and the high-quality reads were compared with the yak reference genome (BosGru3.0, http://ftp.ensembl.org/pub/release-99/fasta/bos_grunniens/), and the resulting BAM files were sorted using the *sort* command of SAMtools **[Li H, et al. 2009]** software (version 1.11) and de-duplicated using the *rmdup* command (no duplicates were marked here, duplicate reads were removed directly). Local recombination of reads and comparison near enhanced indel polymorphisms were performed using Genome Analysis Toolkit (GATK) **[McKenna A, et al. 2010]** software (version 4.0.1). Then the *HaplotypeCaller* command in GATK was used for SNP calling, *CombineGVCFs* command for VCF file merging, *GenotypeGVCFs* command for variant detection, and *VariantFiltration* command for initial filtering, and the variant filtering conditions were set as follows: QD (QualByDepth, variant loci confidence divided by the number of unfiltered non-reference reads) < 2.0 || FS (FisherStrand, Fisher exact test to assess the probability that the current variant is a strand bias, this value is between 0 and 60) > 60.0 || MQ (RMSMappingQuality, square root of the matching quality in all samples) < 40.0 || MQRankSum (MappingQualityRankSumTest, assesses the confidence based on the matching quality of the read of REF and ALT) < 12.5 || ReadPosRankSum (ReadPosRankSumTest, evaluate the variation confidence by the position of the variation in the read, usually the error rate is higher at both ends of the read) < 8.0 || SOR (StrandOddsRati, comprehensive assessment of the likelihood of strand bias) > 3.0. Maker filtering using PLINK **[Purcell S, et al. 2007]** software (version 1.90) with the variant filtering conditions were set as follows: maf (minor allele frequency) 0.05, max-missing(Maximum deletion rate of genotype) 0.05, hwe (deviations from hardy–weinberg equilibrium) 1e6.

### 2.3 Population structure

The NJ-tree was constructed using the P distance matrix calculated by VCF2Dis **[Sun X, et al. 2020]** software (version 1.46), Tree beautification is done in the online site iTol (https://itol.embl.de/), and the principal component analysis (PCA) was performed and plotted using the Genome Associated Prediction Integrated Tool (GAPIT) **[Wang J, et al. 2021]** package in R. The population clustering analysis was done by Admixture **[Chen J, et al. 2021]** software (version 1.30), and the kinship and differentiation between samples from different regions were viewed by joint analysis, and linkage disequilibrium decay analysis was performed using PopLDdecay (version 3.41) **[Zhang C, et al. 2019]**.

### 2.4 Association study

Genome-wide association analysis was performed using a compressed mixed linear model (CMLM) **[Zhang Z, et al. 2010]** in GAPIT (version 3.0) software, where PCA and kinship matrix were added as covariates and *P-values* were corrected with *Bonferroni* and cutoff was set to 0.05/SNP (total number of markers).

The general expressions of CMLM are consistent with mixed linear model (MLM):

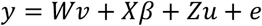

where, *y* is the phenotypic vector (n×1), *v* is the corresponding coefficient vector, which is the non-marker effect among the unknown fixed effects, and *β* is the marker effect among the unknown fixed effects. *W* is the covariate design matrix of vector *v*, *X*is the marker genotype vector of *β*; *u* is the random effect vector of individuals and obeys *u*~*N*(0, *KV_g_*), which *K* is the n×n kinship genetic matrix constructed using the G matrix, *V_g_* is the additive genetic variance; *e* is the random residual and obeys *e*~*N*(0, *IV_e_*), which *I* is the n×n effect matrix, *V_e_* is the residual component; and *Z* is the random effect matrix of *u*.

### 2.5 Population index building

The kinship in the model was calculated with all markers, and combining test markers with kinship in MLM will lead to confusion between test markers and the genetic effects of individuals defined by kinship. In CMLM, Zhang et al. **[Zhang Z, et al. 2010]** introduced a variable called compress group, which clusters individuals with closer kinship into groups and uses the kinship between groups instead of the kinship between individuals and individuals for the operation. In this study, the geographical relationships of individual yaks were artificially classified into groups, and a variable (named *compress_z*) was added to the GAPIT built-in function *GAPIT.Compress.R* to store the real geographical groupings, by providing the groupings in CMLM in advance instead of the compressed groupings being estimated. We name this approach the provided compressed mixed linear model (pCMLM). The specific parameter *compress_z* under this method is marked with the number 1 to be identified as the same group of objects, and the output parameter *group.membership* is marked with the same number to be the same group. The user can add CMLM to GAPIT by providing a real or artificially defined grouping file for the operation, which has been uploaded to github (https://github.com/liu-xinrui/GWAS).

### 2.6 Identification of candidate genes

Based on the physical location of the target trait association loci on the yak reference genome and combined with the LD decay distance of the yak genome (~20kb), associated genes were screened on both sides of the SNP loci. If the gene on the reference genome has only Ensembl ID, the sequence of the gene was extracted for BALST comparison using the wild yak reference genome (BosGru_v2.0, http://ftp.ensembl.org/pub/release-99/fasta/bos_mutus/) for functional analysis. For multiple significant SNP loci, haploid block mapping was performed using LDBlockShow **[Dong S S, et al. 2021]** (version 1.40) for all SNPs within 100kb upstream and downstream of the leader SNP, and association analysis was performed for leader SNPs within a block with phenotypic values on each individual.

## 3 Result

### 3.1 Phenotypic distribution

Here we examined a total of 94 adult yaks for five body sizes traits and summarized the descriptive statistics (mean, variance, maximum, minimum, and coefficient of variation) for different body sizes traits. As shown in Table 1, Most of the recorded body heights ranged from 100 to 205 cm. The mean body height of this yak population was 118.2 cm, body slope length was 138.6 cm, body weight was 284.9 kg, chest circumference was 168.8 cm, and tube circumference was 17.24 cm. all body size data of wild yaks were higher than those of domestic yaks in all regions, and all body size data of wild yaks were maximum. This is due to the fact that yaks eat irregular food and encounter greater external risks, the more natural selection pressure they face, the smaller and less competitive individuals are eliminated. The domestic yak is actually domesticated from the wild yak and is much less wild than the wild yak. Among them, the body size trait of yaks in Tibetan region is in the upper level among domestic yaks.

### 3.2 SNP calling and population structure

A total of 47.15 million markers, including SNPs, Indels and other variants, were detected using the BWA-SAMtools-GATK pipeline **[Wang J, et al. 2022]** program with default parameters. 3,584,464 SNPs remained after filtering by GATK and PLINK software, distributed on average on 29 autosomes and 1 X chromosome. At the 1Mb window size, most markers density was around 2 kbp (Figure 1A). In addition, most individuals’ and SNP markers’ heterozygosity was low (Figure 1B, Figure 1C). These results indicated that the distribution of genetic markers in this study was consistent on chromosomes and at different positions.

**Figure 1.**
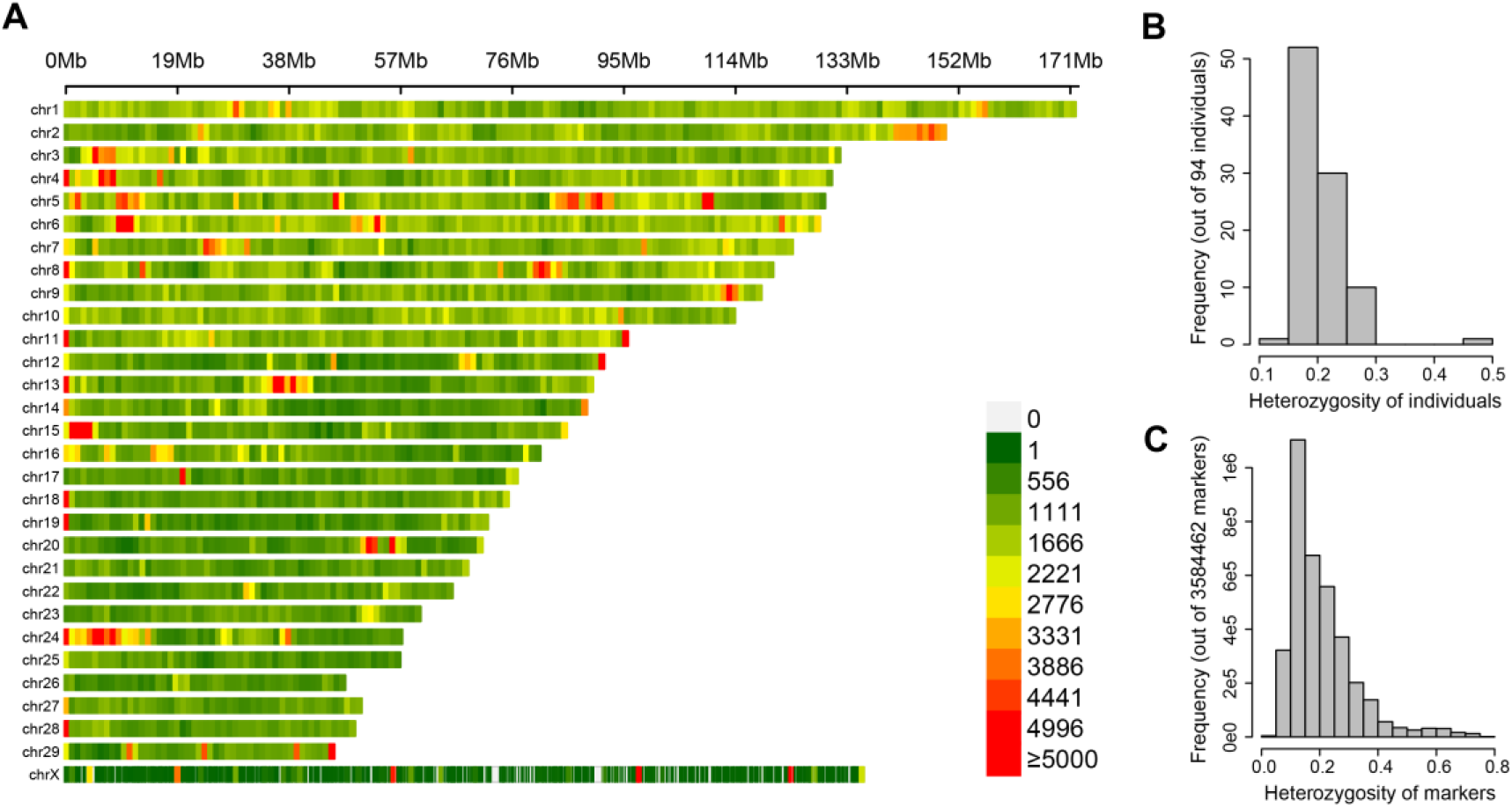
The marker distribution in the whole genome and frequency of heterozygosity. There were 3,584,464 SNP markers in 29 autosomes and one X chromosome. (A) SNP marker density represented by physics position in chromosomes, the more the color leans towards red, the higher the density of markers in the window, each color block indicates the number of SNPs within 1Mb windows size; (B) Histograms of heterozygosity frequencies for all 94 individuals; (C) Histograms of heterozygous frequencies of all SNP markers.

In this study, all yak populations were consisted as multiple yak subspecies in the Qinghai-Tibetan Plateau region of China (Figure. 2A). This sample has multiple breeds and a complex population structure. To analyze the population structure of 94 yaks, we performed principal component analysis, population stratification and NJ-tree analysis on seven subgroups (including 6 regions and wild yak groups) of yaks using 3.58 million high-quality SNP data obtained by filtering. The PCA clustering scatters plot showed that the population structure of this yak population was weak, except for the Tibetan region, which could be roughly clustered into one group, and the other six subgroups were in a mixed state and could not distinguish the population structure (Figure. 2B), and the genetic variance contribution explained by the first 2 principal components was 2.85% and 1.82%, respectively (Figure S1). The NJ-tree clustering results demonstrated approximately the same population structure as PCA (Figure. 2C). The best K value for Admixture population structure analysis was 1, suggesting that the group was closely related and the population structure was weak. Therefore, K was set to 7 in this study, in which Tibetan varieties were clustered into one category (Figure 2E). The linkage disequilibrium analysis showed that the breed-based yak populations had faster decay of linkage disequilibrium and lower LD levels. The most immediate decay was in Tibetan breeds, followed by Sichuan, Qinghai, Gansu, Xinjiang, wild yak and Yunnan.

**Figure.2.**
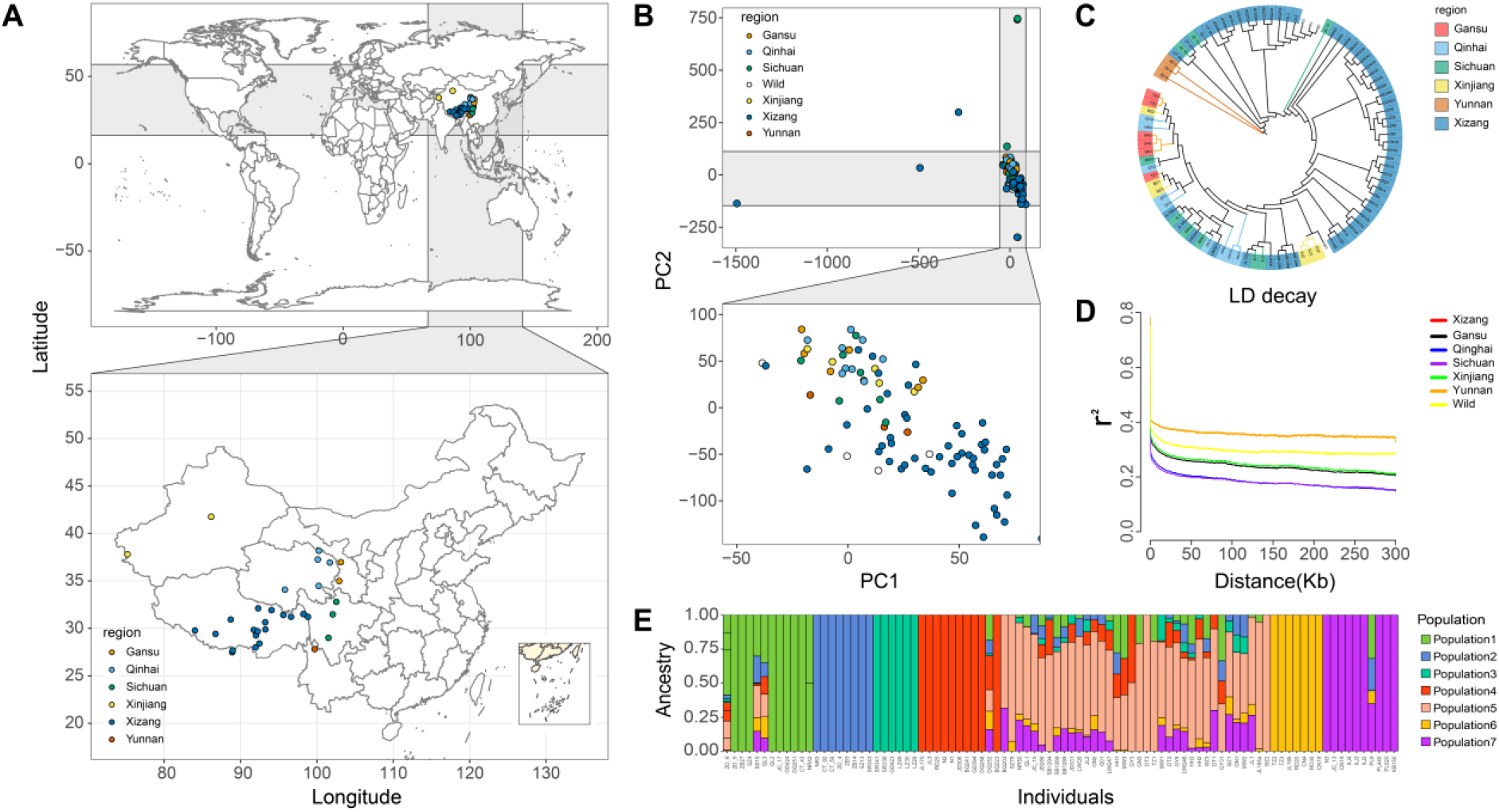
The location distribution of samples collection and population stratification. A total of 31 yak breeds (including one wild yak breed) and 3584463 high-quality SNP markers were collected from 94 individuals. The distribution map of the breeds was drawn using the latitude and longitude of the sampling sites **(A)**. Population structure explained by PCA using all SNP markers **(B)**; All SNP markers were used to generate neighbor-joining (NJ) trees of 94 yaks by VCF2Dis software **(C)**; All SNP markers were used for LD decay analysis of cultivars from seven different provinces and regions (D). All SNP markers were used to cluster the cultivars from 7 different provinces by Admixture software. Different colors are used to indicate different provinces and regions in the figure (**E**).

### 3.3 GWAS and Candidate Genes

The CMLM model was used to estimate the genetic potency of body sizes traits and their statistical power, and two statistical strategies were constructed. The first statistical strategy is the CMLM using a priori population clustering parameters, being named pCMLM group, the second is a tight grouping constructed by splitting all individual kinship matrices into several inter-group kinship matrices using CMLM, which is named CMLM group. The -2LL (Twice negative log likelihood) of the two strategies were compared for BH traits with significant loci detected, with the pCMLM (561.58) outperforming the CMLM (509.73). Genotypic and phenotypic data of 94 individual yaks were analyzed using GAPIT software with PCA and kinship as fixed effects. Manhattan and QQ-plots are shown in Figure. 3. After Bonferroni correction, six SNPs were found to pass the 5% threshold line (*P-value* < 1.39 × 10-8) and correlated with BH only in the pCMLM (Figure 3A), where no significant loci associated with body sizes were detected in any of the CMLM (Figure S2-6). The QQ-plot of BH in the pCMLM showed that the scatter drifts gradually above the diagonal after the expectation value equals 2, indicating that the population was subjected to natural selection revealed that the effect of loci exceeded the random mark, and the analytical model was reasonable. The likelihood value used to determine the best compression ratio in traditional CMLM is not significant. CMLM does not catch the best clustered groups, which is reflected in the low statistical power of the association results (Figure 3B). QQ plots of the traditional CMLM with BH show that most points lie below the diagonal (Figure 3C), which indicates that the observed P-value for most loci is less than the expected value and that the model overcorrects for this group. The significant SNPs detected in the pCMLM were rs769892 on chromosome 4, rs2659279 and rs2659285 on chromosome 13, rs310769 and rs477265 on chromosome 2, and rs2910497 on chromosome 15, which were located at 4,883,046 bp, 59,118,279 bp, 59,119,427 bp, 16,165,590 bp, 57,505,237 bp, and 129,838,648 bp, respectively. The significant SNPs with their associated candidate genes did not have annotation information in the 100kb linkage disequilibrium interval information each upstream and downstream. Therefore, their FASTA sequences were extracted and matched to the wild yak reference genome, and all of them were compared to the associated genes except for the rs769892 locus with the highest statistical correlation with the body height trait, which failed to match the associated genes (Table 3).

**Figure.3.**
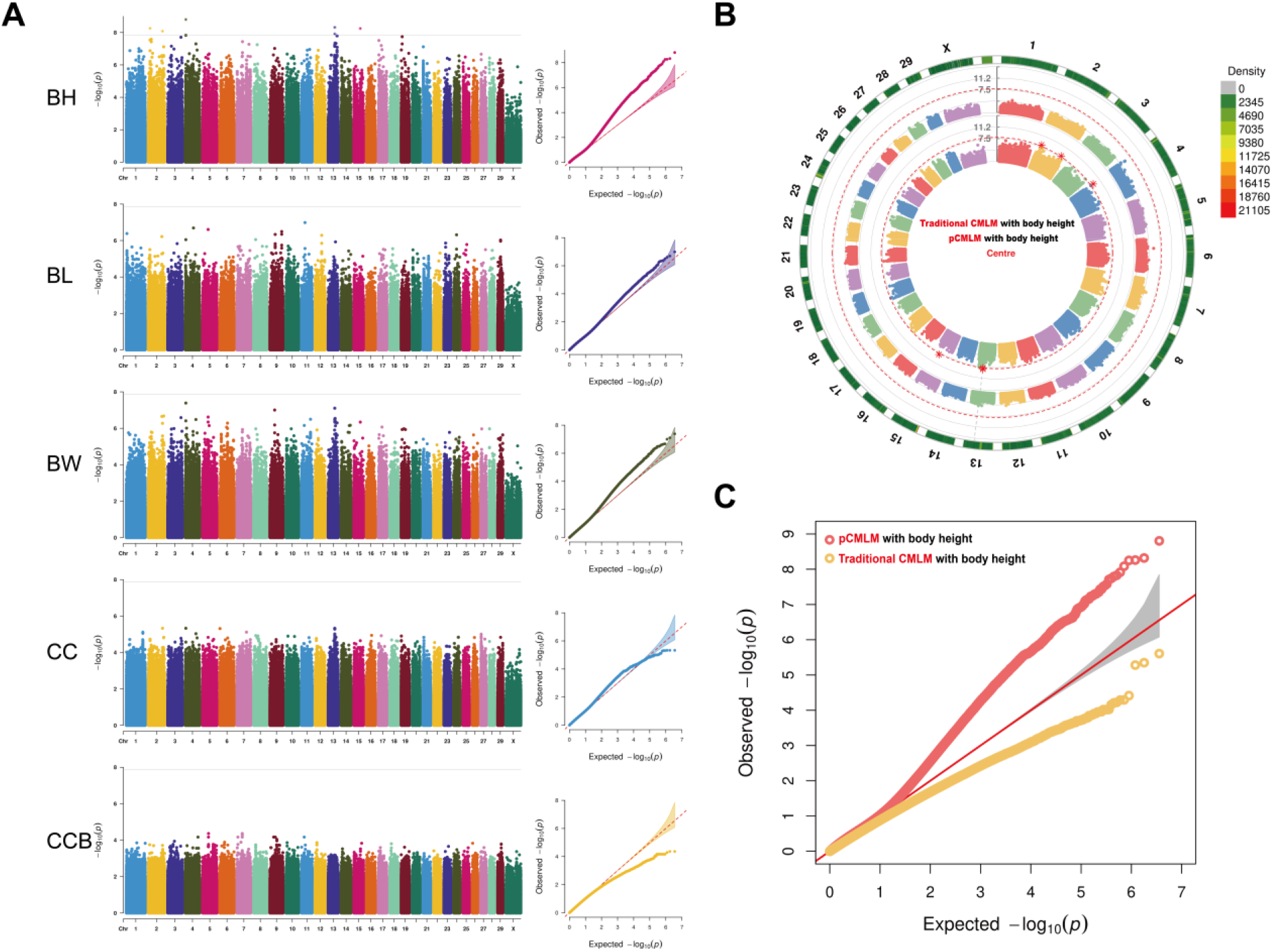
Manhattan and quantile-quantile plots of the *p-values* for the genome-wide association study of body height (BH), body length (BL), body weight (BW), chest circumference (CC) and circumference of cannon bone (CCB) of yaks based on the pCMLM method, the horizontal line of significance threshold (*p-value* < 1.39 × 10^-8^) was used to distinguish significantly associated loci, and the different colors to distinguish different chromosomes (A); Circular Manhattan (B) and quantile-quantile plots (C) of the optimized pCMLM method and the conventional CMLM method in detecting body height traits in yaks, where the inner ring is pCMLM, the outer ring is conventional CMLM, and the outermost ring indicates the labeling density of this chromosome.

**Table 3.**
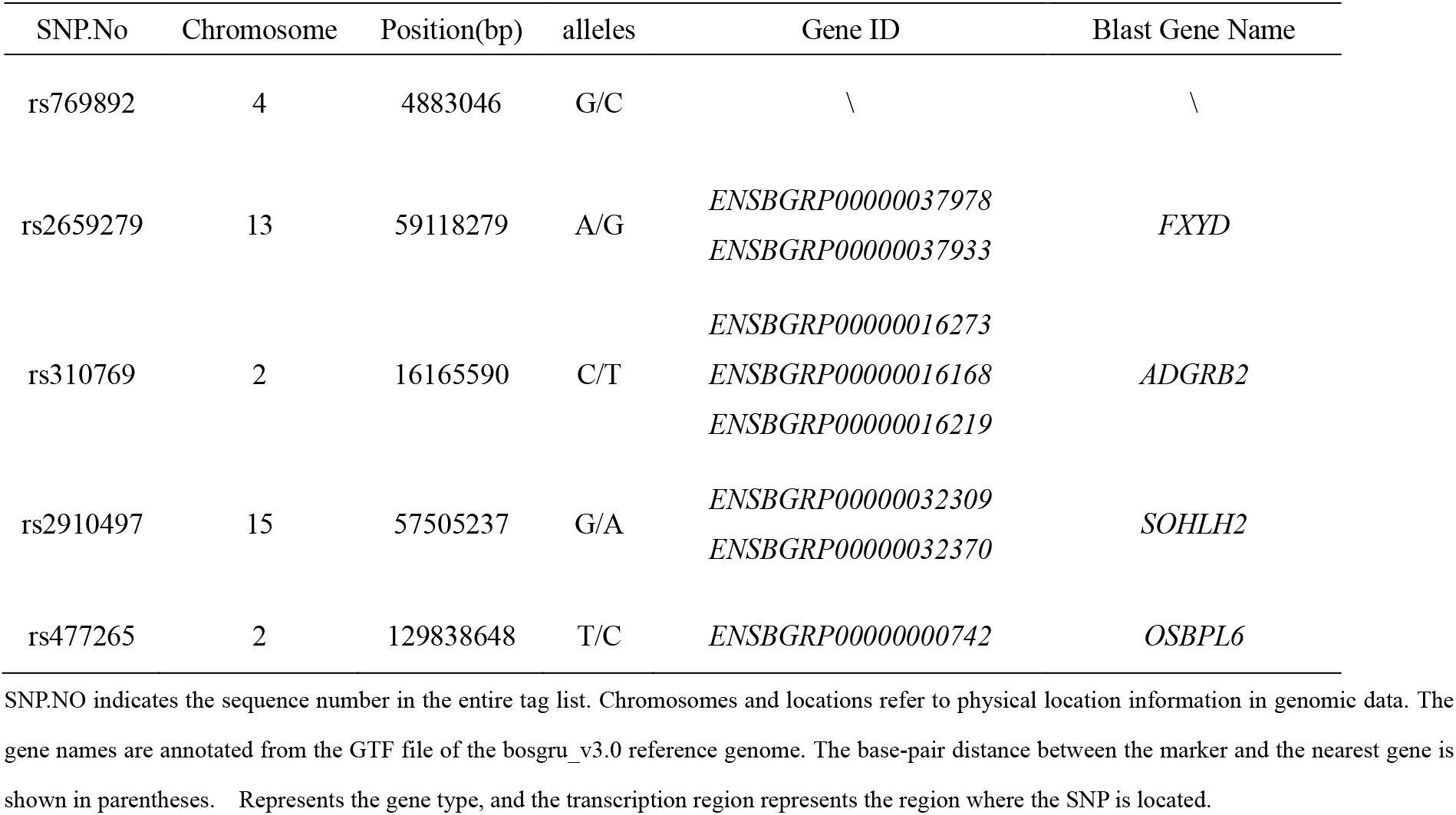
Relevant significant SNP information of candidate genes in the 200kb region.

### 3.4 Genotype correlation in the LD Block

To further identify candidate genes for BH traits, we evaluated the allelic effects associated with BH, and some SNPs showed significant allelic effects on BH, indicating that candidate genes for BH may be associated with these SNPs. The results of haplotype analysis of 100 kb before and after the Leader SNP showed that there was linkage disequilibrium in chr15-rs2910497-57505237 (chr: chromosome of the SNP; rs: SNP numbering on the genome; the third digit indicates the relative physical position of the chromosome in which it is located), the nearby LD block region was small, and the SNP was located on the outer side of the transcript. The analysis of allele haplotype based on this SNP showed that only GA and GG genotypes existed in 94 individuals, and no homozygous AA genotype was observed. Yak individuals with allele A showed higher BH effect (Figure 4A). The same situation was also observed for four SNPs, rs2659279 (Figure 4B), rs477625 (Figure 4C), rs310769 (Figure 4D) and rs769892 (Figure 4E), and the four haplotypic alleles with strong additive effects on BH were G, C, T and C, respectively, among the respective haplotypic alleles. There is a strong LD between chr13-rs2659279-59118279 and chr4-rs769892-4883046 two SNPs in a LD block, but the block segment is small. Among them, rs2659279 and rs2659285 on chromosome 13 have strong linkage disequilibrium, and the two SNPs are physically close to each other, and rs2659279 is used as a benchmark when screening candidate genes. Except for rs769892, the other four SNPs with statistical significance were located near the intron region where at least one unknown transcript existed, and the sequences of the physically closest transcripts were compared by blast. The results showed that the transcript near rs2659279 was matched to *FXYD* gene, rs310769 to *ADGRB2,* rs2910497 to *SOHLH2,* and rs477265 to *OSBPL6* (Table 3).

**Figure.4.**
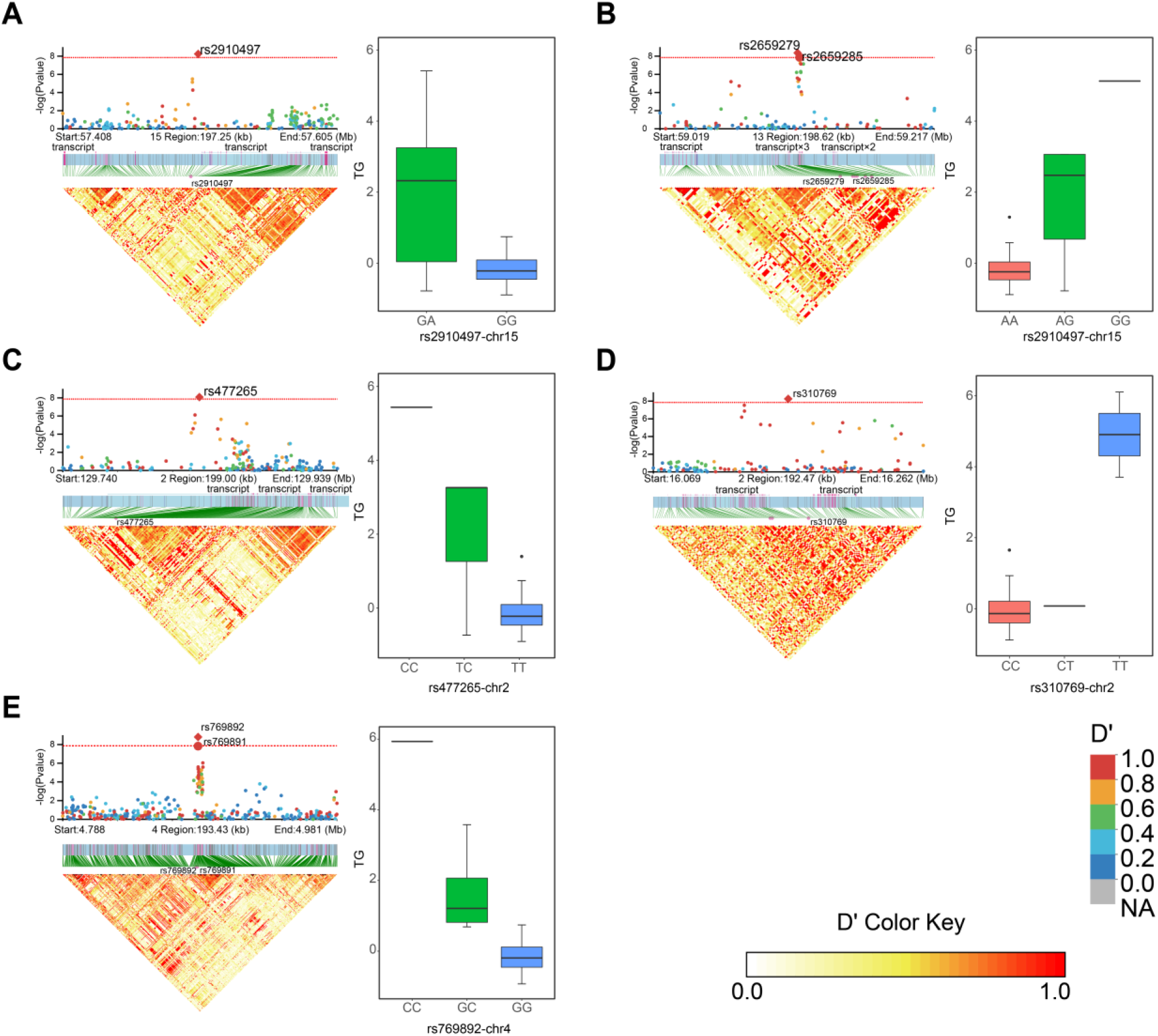
Local Manhattan plot of 200 kb near each significant leader SNP (top left), the structure of the transcript in this region (middle left) and the LD heat map (bottom left). Phenotype distribution among the genotypes of all leader SNPs on the right side. Where the red horizontal line indicates the threshold (*p-value* < 1.39 × 10^-8^) of GWAS. Multiple transcripts are abbreviated in the figure. The LD heat map distinguishes the strength of association with different colors, and the redder the color, the stronger the association. The genotype-phenotype association plots of SNPs distinguishes different genotypes in pink, green and blue. rs2910497 of chromosome 15 (A), rs2659279 of chromosome 13 (B), rs477265 of chromosome 2 (C), rs310769 of chromosome 2 (D), and rs769892 of chromosome 4 (E). Only two genotypes, GA, and GG, were observed for rs2910497-chr15 in all individuals, but heterozygous genotypes were observed for the other four leader SNPs.

## 4 Discussion

Whether it is marker-assisted selection (MAS), genome selection (GS) or gene editing (GE) technology, it is necessary to accurately locate the genetic markers of target traits. GWAS, an effective method for finding molecular makers affecting essential economic traits, is commonly used to directly associate complex trait phenotypes with genome-wide single nucleotide polymorphisms in a large number of individuals in the same population **[Breeze C E, et al. 2022]**. It is widely used to locate human disease loci **[Tam V, et al. 2019]** and develop molecular markers **[Tian D, et al. 2020]**, as well as to aid in plant and animal breeding selection **[Jeong S, et al. 2020]**. However, GWAS analysis based on multi-population and multi-variety has not been reported. As a large livestock species in the Qinghai-Tibetan plateau region of China, yak has many breeds and local breed resources due to special geographical reasons. The population structure of multiple breeds is often complex. There will be pronounced population stratifica tion, and the population structure is an essential factor affecting the accuracy of association results, which often leads to false-positive association results **[Bouaziz M, et al. 2011]**.

Variation in body size traits is mainly based on distant crosses of different yak breeds. In particular, current domestic yak production performance has decreased and herders are using a large number of wild yaks with more superior traits for breeding. This wild-blooded yak causes mutation loci associated with wild yak body size traits to be exchanged among domesticated yaks. Therefore, yak populations are more complex throughout the whole Tibetan Plateau region. None of the 31 yak breed resources in this study showed significant population clustering in multiple cluster analyses, with only the Tibetan breeds having a more similar genetic structure due to their domesticated origins in Tibet **[Chai Z X, et al. 2020]**. Such populations with complex population structure are not conducive to statistical analysis by traditional CMLM. The grouping process of the CMLM, which assigns individuals with similar characteristics to the same group, uses the elements in the kinship matrix as a similarity measure **[Li M, et al. 2014]**, replacing the kinship between individuals and individuals with the kinship between groups and groups **[Zhang Z, et al. 2010]**. The genetic principle utilizes intra-group balance, thus reducing the variance of the model’s residual part, which will improve statistical power for GWAS. The difference between pCMLM and CMLM is that pCMLM provides clustering relationship of species or taxa in advance, and the kinship is compressed directly. In contrast, CMLM needs to filter the best compression ratio by likelihood value, however, sometimes the likelihood values between different compression ratios are not very significant. The above situation would cause CMLM to use individuals to represent groups and kinship without any compression, thus making CMLM back to MLM. Group structure or population origin in yak populations usually also implies similarity in nutritional level and growth environment **[Guo N, et al. 2021]**, so using these factors to force the CMLM to be compressed is beneficial for detecting candidate genetic markers. However, the complex structure will make the statistical power of the CMLM closer to that of the standard MLM. Therefore, when the traditional CMLM cannot catch the optimal clustered groups, our proposed pCMLM can also reduce the variance of the residuals in the model by providing the real kinship and group structure. This can provide us with more adequate association results.

Therefore, GWAS results based on pCMLM identified six SNP loci that were statistically significant in association with BH. Based on the annotation information provided by the 100 kb upstream and downstream of the yak reference genome, we obtained relevant annotation information on only four SNPs, but these contained only Ensembl ID. Probably because the yak reference genome is not yet complete, the sequences of these genes were compared to the wild yak reference genome by BALST tool, and the candidate genes showed that rs2659279 (chr13-59118279) was associated with *FXYD6*(domain containing ion transport regulator 6), rs2910497 (chr15-57505237) was associated with *SOHLH2(spermatogenesis* and oogenesis specific basic helix-loop-helix 2), and rs310769 (chr2-16165590) associated with *ADGRB2(adhesion* G protein-coupled receptor B2), and rs477265 (chr2-129838648) associated with *OSBPL6*(adhesion G protein-coupled receptor B2). Among these genes, the *FXYD6* gene is a member of the *FXYD* family encoding a transmembrane protein, a specific protein encoding the hippocampus phosphate that has been shown in humans to be involved in mediating the Na/K ion pump **[Gao Q, et al. 2014]**, *FXYD6* significantly accelerates Na^+^ deactivation and Na^+^ pump conversion rates **[Garty H, et al. 2006]** and alters the selectivity of the intracellular ion pump **[Meyer D J, et al. 2020]**. The *SOHLH2* gene belongs to the *b-HLH* (Basic helix-loop-helix transcription factor) family, which encodes a testis-specific transcription factor essential for spermatogenesis, oogenesis, and folliculogenesis **[Ballow D J, et al. 2006]**. The *b-HLH* family is involved in numerous biological processes in the organism, including cell differentiation, cell cycle arrest, and apoptosis **[Sun H, et al. 2007]**. The *SOHLH2* gene was also shown to play an important regulatory role in the reproductive gonadal axis, pituitary, hypothalamus, ovary and testis in buffalo **[Zhun P, et al. 2016]**, pig **[Liu K, et al. 2016]** and mouse **[Ballow D J, et al. 2006; Park M, et al. 2016; Wang Z, et al. 2020]**. The *ADGRB2 gene* acts as a transcriptional repressor through GA-binding protein, regulates vascular endothelial growth factor and is significantly associated with its growth traits in grouper **[Yu H, et al. 2018]**. The *OSBPL6* gene is a member of the family encoding hydroxysteroid-binding protein (OSBP), an intracellular lipid receptor **[Herold C, et al. 2016]**. The *OSBPL6* gene contributes to the maintenance of cholesterol homeostasis in vivo by regulating cholesterol transport in humans through miR-33 and miR-27b **[Ouimet et al., 2016],** and in studies of *OSBPL6* in juvenile DePaul dwarf horses, it was shown that *OSBPL6* is an essential factor affecting body height in DePaul dwarf horses **[Pan J, et al. 2019]** by showing variable splicing of ES type in the pituitary gland, and ES variable splicing causes GH1 third exon jumping resulting in a 17.5 kD GH isoform, which is an essential factor contributing to height defects in patients with autosomal dominant growth hormone deficiency (type II) **[Alatzoglou K S, et al. 2015]**, and the gene may be associated with multiple epiphyseal dysplasias **[Jeong C, et al. 2014]**. Therefore, it is hypothesized that two genes, *OSBPL6* and *ADGRB2*, are the most likely candidates to affect body height traits in yaks. However, whether and how the studied localized genes affect body height and size traits in yaks needs to be further explored, providing new directions and ideas for later validation studies.

## 5 Conclusion

In conclusion, the pCMLM, which provides a previous population information, can achieve better statistical results when the traditional CMLM cannot provide effective compression grouping by obtaining optimal likelihood values. The GWAS results of pCMLM associated six SNPs with body height and screened four candidate genes, *FXYD6*, *SOHLH2*, *ADGRB2*, and *OSBPL6*, with body height phenotypes. This study will help us to develop better biostatistical model optimization ideas and deeper understanding of the relationship between genes and body height, and the results may provide basic information for quantitative trait gene localization or candidate gene cloning in yak body height formation mechanism.

## Declarations

### Ethics approval and consent to participate

Not applicable

### Consent for publication

Not applicable

### Competing interests

The authors declare that they have no competing interests.

### Funding

This project was partially funded by the Qinghai Science and Technology Program, China (2022-NK-110), Sichuan Science and Technology Program, China (Award #s 2021YJ0269 and 2021YJ0266), the Fundamental Research Funds for the Central Universities, China (Southwest Minzu University, ZYN2022023), and the Program of Chinese National Beef Cattle and Yak Industrial Technology System, China (CARS-37).

### Authors’ contributions

**XL:** Software, Writing first draft of manuscript, Visualization, Testing, Methodology, Supervision, and Validation**. ZC, WP, YK, and JZ:** Writing first draft of manuscript. **JW:** Conceptualization and Manuscript Revision.

## Acknowledgments

We would like to thank Editage (www.editage.cn) for English language editing.

## Notes

https://github.com/liu-xinrui/GWAS

### Competing Interest Statement

The authors have declared no competing interest.

