## Supplemental Table and Figure 1-7 for "pCMLM: Genome Wide Association Study of Body Size Traits in Multiple Regions of Yak Based on the Provided Compressed Mixed Linear Model"

### Supplementary Material

**Supplementary Table.1** Distribution area of yak of different variety resources

| Variety resources | Location (provinces or autonomous) | Sampling area | Altitude (m) | Average annual team. (℃) |
|---|---|---|---|---|
| Zhongba | Tibet | Zhongba county, Shigatse, Tibet autonomous region | 5000 | -4 to 12 |
| Shenza | | Located in central Tibet of Shenza county, which between the Gangdis Mountains and the second largest lake in northern Tibet | 4700 | 0.5 |
| Cona | | Located in the southern tip of the Tibet Autonomous Region and the Himalayas southeast of the Cona county | 4380 | -4 to 8 |
| Sangsang | | Ngamring County, Shigatse, Tibet autonomous region | 4400 to 4600 | 6.5 |
| Sangri | | Sangri County, Lhoka, Tibet autonomous region is located at the southern foot of the Gangdis Mountains and in the middle valley of the Yarlung Tsangpo River. | >4065 | 8 |
| Sibu | | Mozhugongka County, Lhasa City, Tibet Autonomous Region is located in the middle and upper reaches of Lhasa River and the west side of Mila Mountain | 3835 | 2 to 17 |
| Riduo | | Riduo Township, Mozhugongka County, Lhasa City, Tibet Autonomous Region | 4370 | 0.8 |
| Pali | | Yadong County, Shigatse, Tibet Autonomous Region | 3500 | 0 |
| Nierong | | Nyainrong County, Naqu, Tibet Autonomous Region | 4700 | -7 to 7 |
| Longzi | | Lhünzê County, Lhoka, Tibet autonomous region | 3900 | 5.5 |
| Leiwuqi | | Riwoqê County, Qamdo, Tibet Autonomous is located in a branch of the Nyainqêntanglha Shanmai and the west of the Boshula Ridge | 4500 | 2.5 |
| Kangbu | | Kangbu Township, Yadong County, Shigatse, Tibet Autonomous Region | >3500 | - |
| Lijia | | Lhari County, Naqu, Tibet Autonomous Region | 4500 | -6 to 8 |
| Jiangda | | Jomda County, Qamdo, Tibet Autonomous Region | 3650 | 4.5 |
| Gongbujiangda | | Gongbo'gyamda County, Nyingchi, Tibet Autonomous Region | >3600 | 3 to 16 |
| Baqing | | Baqing County is located in the northeast of Tibet Autonomous Region, the east of Naqu District, and the upper reaches of salween | >4500 | -1 |
| Dingqing | | Dênqên County, Qamdo, Tibet Autonomous Region | 3850 | 3.4 |
| Qilian | Qinghai | Qilian County is located in the north of Haibei Tibetan Autonomous Prefecture of Qinghai Province, bordering Gansu Province in the east and north | >2800 | 1 |
| Huanhu | | Gangcha County, Haibei Tibetan Autonomous Prefecture of Qinghai Province, is located in the north of Qinghai Lake | 3300 | -0.6 |
| Gaoyuan | | Tibetan Autonomous Prefecture of Golog, Qinghai Province | 4000 to 5000 | -4 |
| Datong | | Datong Hui and Tu Autonomous County, Xining, Qinghai Province | 2280 to 4622 | 4.9 |
| Maiwa | Sichuan | Hongyuan Country, Tibetan Qiang Autonomous Prefecture of Ngawa, Sichuan Province | 3500 | 2.9 |

| | | | | |
|---|---|---|---|---|
| Jiulong | | Jiulong Country, Tibetan Qiang Autonomous Prefecture of Ngawa, Sichuan Province | 1440 to 6000 | 4.9 |
| Changtai | | Baiyu County, Tibetan Qiang Autonomous Prefecture of Ngawa, Sichuan Province | 3500 | 12.3 |
| Jinchuan | | Jinchuan Country, Tibetan Qiang Autonomous Prefecture of Ngawa, Sichuan Province | 1950 to 5000 | 12.4 |
| Tianzhu | Gansu | Tianzhu Zangzu Autonomous County, Wuwei, Gansu Province | 2040 to 4874 | 3.5 |
| Gannan | | Maqu County is located at the east end of the Qinghai Tibet Plateau and the southwest of Gannan Tibetan Autonomous Prefecture | 3700 | 1.2 |
| Xinjiang | Xinjiang | Tajik Autonomous County of Taxkorgan, Kashgar Prefecture, Xinjiang Autonomous Rgion | 3600 | 3.3 |
| Bazhou | | Korla City, Bayingol Mongolian Autonomous Prefecture, Xinjiang Autonomous Rgion, is Located in the middle of Xinjiang, the southern foot of Tianshan Mountains, and the northeast edge of Tarim Basin | 934 | 11.4 |
| Zhongdian | Yunnan | Zhongdian county in Diqin Tibetan autonomous prefecture | 3450 | 5.5 |
| Wild yak | Qinghai-Tibet Plateau | - | - | - |

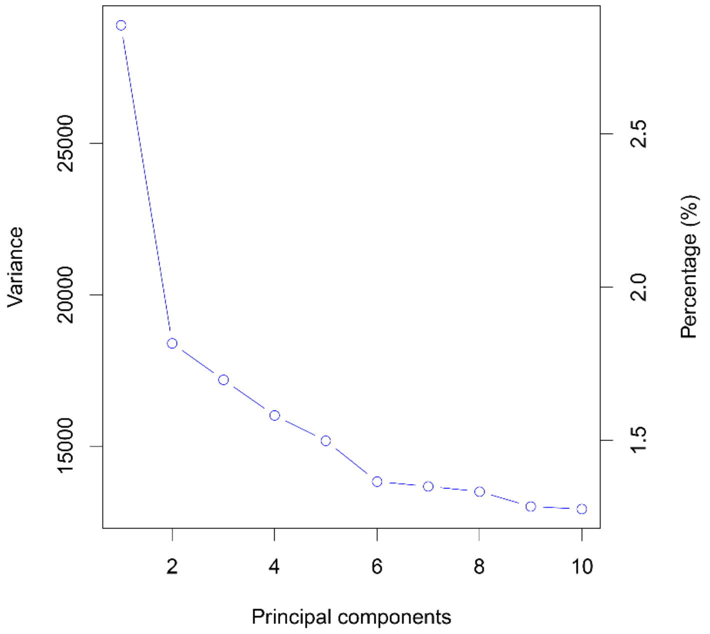


**Supplementary Figure.1** Genetic variance explained and percentage of overall with Principal components of each dimension. The horizontal coordinates indicate the principal components of the different dimensions; The left vertical coordinate indicates the genetic variance explained by each principal component; The right vertical coordinate indicates the percentage of the overall variance accounted for by each principal component.

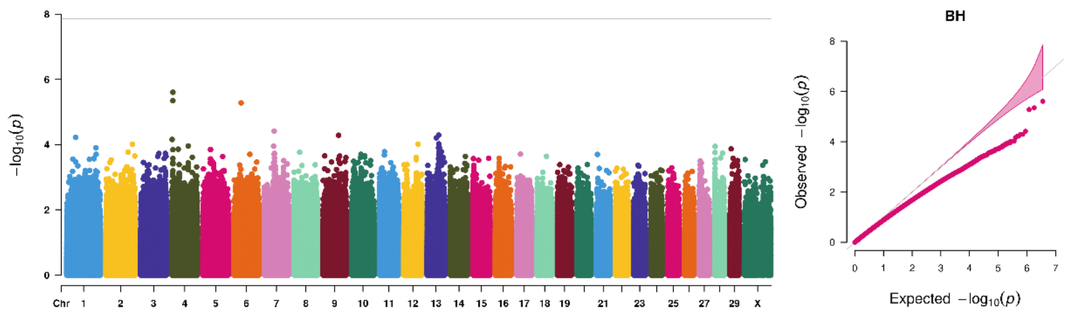


**Supplementary Figure.2** Manhattan (left) and quantile-quantile plots (right) of the *p-values* for the genome-wide association study for body height (BH) of yaks based on the traditional CMLM method, the horizontal line of significance threshold (*p-value* $< 1.39 \times 10^{-8}$) was used to distinguish significantly associated loci, and the different colors to distinguish different chromosomes.

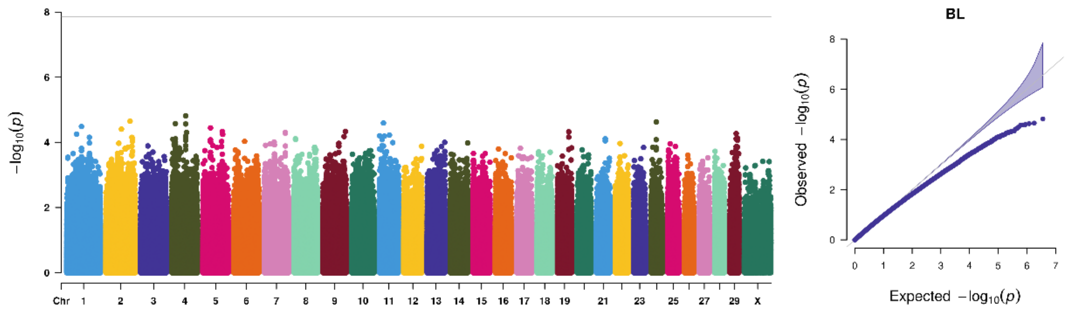


**Supplementary Figure.3** Manhattan (left) and quantile-quantile plots (right) of the *p-values* for the genome-wide association study for body length (BL) of yaks based on the traditional CMLM method, the horizontal line of significance threshold (*p-value* $< 1.39 \times 10^{-8}$) was used to distinguish significantly associated loci, and the different colors to distinguish different chromosomes.

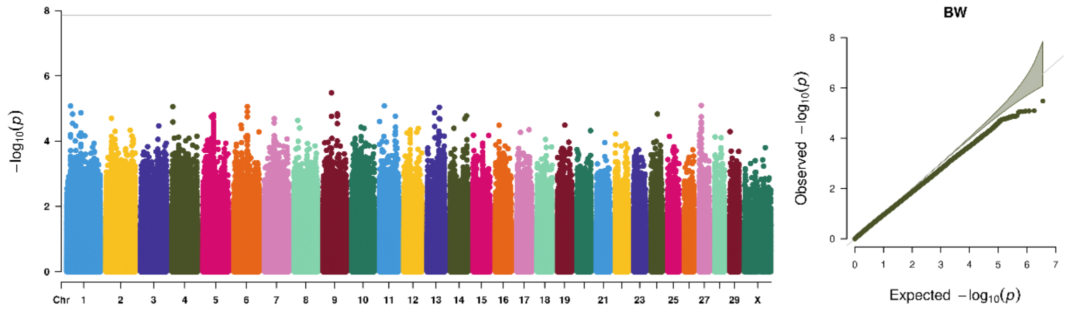


**Supplementary Figure.4** Manhattan (left) and quantile-quantile plots (right) of the *p-values* for the genome-wide association study for body weight (BW) of yaks based on the traditional CMLM method, the horizontal line of significance threshold (*p-value* $< 1.39 \times 10^{-8}$) was used to distinguish significantly associated loci, and the different colors to distinguish different chromosomes.

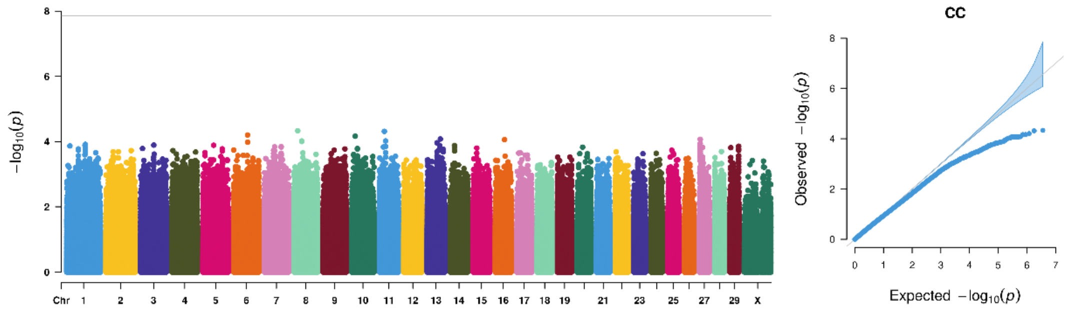


**Supplementary Figure.5** Manhattan (left) and quantile-quantile plots (right) of the *p-values* for the genome-wide association study for chest circumference (CC) of yaks based on the traditional CMLM method, the horizontal line of significance threshold (*p-value* $< 1.39 \times 10^{-8}$) was used to distinguish significantly associated loci, and the different colors to distinguish different chromosomes.

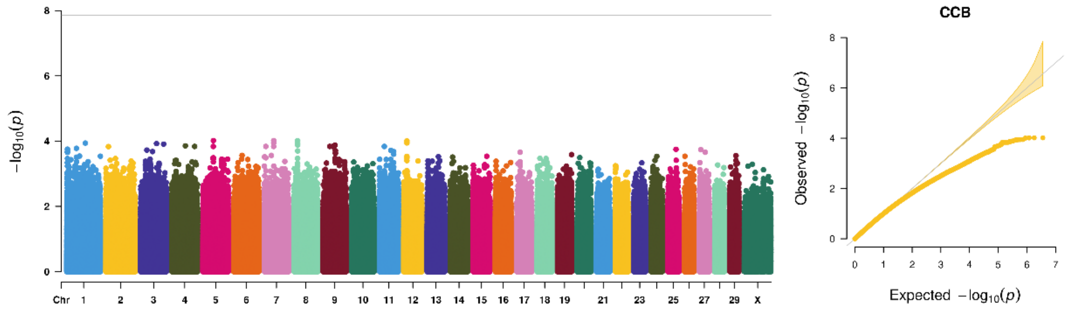


**Supplementary Figure.6** Manhattan (left) and quantile-quantile plots (right) of the *p-values* for the genome-wide association study for circumference of cannon bone (CCB) of yaks based on the traditional CMLM method, the horizontal line of significance threshold (*p-value* $< 1.39 \times 10^{-8}$) was used to distinguish significantly associated loci, and the different colors to distinguish different chromosomes.
